# MeShClust: an intelligent tool for clustering DNA sequences

**DOI:** 10.1101/207720

**Authors:** Benjamin T. James, Brian B. Luczak, Hani Z. Girgis

**Affiliations:** Bioinformatics Toolsmith Laboratory, Tandy School of Computer Science, University of Tulsa, 800 South Tucker Drive, Tulsa, OK 74104, USA; Mathematics Department, University of Tulsa, 800 South Tucker Drive, Tulsa, OK 74104, USA

## Abstract

Sequence clustering is a fundamental step in analyzing DNA sequences. Widely-used software tools for sequence clustering utilize greedy approaches that are not guaranteed to produce the best results. These tools are sensitive to one parameter that determines the similarity among sequences in a cluster. Often times, a biologist may not know the exact sequence similarity. Therefore, clusters produced by these tools do not likely match the real clusters comprising the data if the provided parameter is inaccurate. To overcome this limitation, we adapted the mean shift algorithm, an unsupervised machine-learning algorithm, which has been used successfully thousands of times in fields such as image processing and computer vision. The theory behind the mean shift algorithm, unlike the greedy approaches, guarantees convergence to the modes, e.g. cluster centers. Here we describe the first application of the mean shift algorithm to clustering DNA sequences. MeShClust is one of few applications of the mean shift algorithm in bioinformatics. Further, we applied supervised machine learning to predict the identity score produced by global alignment using alignment-free methods. We demonstrate MeShClust’s ability to cluster DNA sequences with high accuracy even when the sequence similarity parameter provided by the user is not very accurate.

## INTRODUCTION

Clustering nucleotide sequences is an essential step in analyzing biological data. Pioneering sequence clustering tools have been proposed for reducing redundancy and correcting errors in the next-generation sequencing data (1–4) and for assembling genomes de-novo (5–7). Sequence clustering tools were also proposed for barcode error correction (8) and for taxonomic profiling (9). In addition, d2 cluster (10), CD-HIT (3, 11), UCLUST (12), DNACLUST (9), mBKM (13), and d2-vlmc (14) are general-purpose sequence clustering tools. These tools are applied to clustering gene sequences, expressed sequence tags, RNA, and reducing a set of sequences to a non-redundant group of sequences.

Some of the most widely-used tools for sequence clustering, such as CD-HIT and UCLUST, depend on greedy algorithms, which are not guaranteed to find the optimal solution. Given the importance of sequence clustering in the field of computational biology, we propose a much more advanced approach. The mean shift algorithm is a general-purpose optimization technique (15), which has been widely applied in image processing and computer vision (16–18). Unlike the related greedy approaches, the mean shift algorithm is “guaranteed” to converge to local optimal points, e.g. a center of a cluster. Although this algorithm has been applied successfully thousands of times in other fields, it has been applied only few times in the field of bioinformatics (19–21). Here, we propose novel software, MeShClust, utilizing the mean shift algorithm in clustering nucleotide sequences. Further, our adaptation of the algorithm utilizes a novel classifier to predict the identity score using four alignment-free sequence similarity measures.

In practice, the underlying sequence similarity that separates clusters is not known; therefore a biologist may have to guess an identity score to provide to the clustering tool. If wrong, this guessed score limits the quality of the predicted clusters remarkably. For example, if the provided identity score were higher than the true identity score, a tool would produce smaller clusters; if it were much lower, a tool would produce larger clusters. In both situations, the predicted clusters do not match the real clusters.

Further, the related tools are based on greedy algorithms, in which the selection of the sequence representing the center of a cluster is not necessarily optimal. In these algorithms, a sequence that does not belong to any cluster is considered the center of a new cluster. Once a center is selected, it does not change. To illustrate, if the center sequence is at the periphery of the real cluster, then the predicted cluster is very likely to be a partial cluster. Because the core of MeShClust is the mean shift algorithm, it overcomes these two limitations. Specifically, MeShClust is flexible and is capable of correcting the provided identity score to a great extent. In addition, the sequence representing a cluster *does* change, moving toward the true center of the cluster. Thus, MeShClust provides a stable clustering algorithm that is not very sensitive to the sequence similarity parameter and provides greater accuracy than its counterparts.

## MATERIALS AND METHODS

### Overview

Algorithms 1 and 2 give an overview of the methods underlying our software tool, MeShClust. The software consists of these two components: (i) a classifier and (ii) the mean shift algorithm.

The classifier predicts whether or not two sequences are similar to each other. The similarity is measured as the identity score based on the global alignment of the two sequences (22, 23). Sequences are represented as histograms of counts of short words in the sequences. The classifier predicts the identity score due to global alignment (22, 23) by calculating a weighted sum consisting of few alignment-free similarity measures using a General Linear Model (GLM).

**Algorithm 1** An overview of the algorithm implemented in MeShClust

**Input:** A set of *n* nucleotide sequences *S* = {*s*_1_, *s*_2_, …, *s*_*n*_} sorted by decreasing length

**Output:** Clusters of sequences and their respective centers

If the identity is above 60%, train the classifier using a subset of *S* to recognize similar sequences

*center_cur_* ← *s*_1_

*cluster_cur_* ←{*s*_1_}

**while** *S* is nonempty **do**

*G←* all sequences from *S* close to *center_cur_*

*S←S‒G*

**if** *G* is nonempty **then**

*cluster_cur_* ← *cluster_cur_* ∪*G*

Run *MeanShift*(*center_cur_, cluster_cur_*) to update *center_cur_* (Algorithm 2)

**else**

Add *center_cur_* to *Centers*

Add *cluster_cur_* to *Clusters*

*center_cur_ ←* the closest sequence in *S* to the Old *center_cur_* according to the Czekanowski similarity(Equation **4**) or the identity score if the alignment algorithm is used instead of the classifier

*center_cur_* ←{*cluster_cur_*}

**end if**

**end while**

**for** i = 1 to *num_ iter* **do**

**for all** *center_j_* ∈ *Centers, cluster_j_* ∈ *Clusters* **do**

Run the mean shift to update *center_j_* using sequences in *cluster_j_* along with neighboring clusters (Algorithm 2)

**end for**

**for** *j* = 1 to *|Centers|* **do**

**for all** *center_k_ ∈ Centers* close to *center_j_* **do**

Merge centers *center_j_* and *center_k_* if the sequences representing *center_j_* and *center_k_* are similar

**end for**

**end for**

**end for**

**Algorithm 2** Sequence clustering using the mean shift

**Input:** The current center, *center_cur_*, of a cluster, and a set of points, *X*

**Output:** The closest point in *X* to the new center

Calculate the new center, *center_new_*, using the current center, *center_cur_*, according to Equation **1**

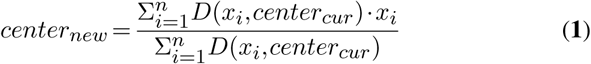

Here, *D* is the classifier or the alignment algorithm if the classifier is not used (Equation **2**)

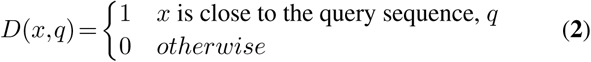

Next, the closest point *p∈ X* to *center*_*new*_ is found using the Czekanowski similarity (Equation **4**) or the identity score if the classifier is not used, and *p* is returned as the new center

A novel adaptation of the mean shift algorithm (15) is the core of the second component. Similar to the classifier, the mean shift algorithm processes the histograms of the input sequences. The mean shift is an iterative, gradient-ascent algorithm that is capable of finding local optimal points. In this adaptation of the algorithm, a local maximum represents the center of a cluster of sequences. In each iteration, a center is recalculated as the weighted *mean* of histograms. This weighted mean is calculated only from the sequences that are similar to the center of a cluster. Similar sequences are determined by the classifier or, if the identity score is below 60%, they are determined by the alignment algorithm. Once updated, a center will *shift* toward a local maximum. As these centers move, some of them converge to the same local maximum; therefore, the algorithm merges them. For this reason, the user does not need to specify the number of centers as opposed to other clustering algorithms such as k-means based applications. Once the algorithm converges, sequences that contributed to the calculation of a center are considered members of its cluster. Supplementary Data Sets 1–3 contains the source code and the executables of MeShClust.

Next, we give the details of each step of the algorithm. First, we describe how a sequence is represented as a k-mer histogram. Second, the details of the classifier are given. In the third step, the initial clusters are formed. We illustrate the construction of the final clusters in the forth step.

### Representing a sequence as a histogram of k-mers

A sequence consists of the nucleotides: A, C, G, and T (or U). A k-mer is a short subsequence of length k. For example, AAA, AAC, AAT, and AAG are tri-mers. To construct a histogram from a sequence, A, C, G, and T are converted to 0, 1, 2, and 3, so a k-mer is built as a quaternary number of *k* digits. Horner’s rule can be used for calculating the quaternary numbers of a long sequence efficiently (24). The count of a k-mer in the histogram is initialized to 1 instead of 0; these pseudocounts are needed to allow events that “seem” impossible to be able to happen (25). For example, k-mers that are absent from one sequence could be present in another. Pseudocounts are important while calculating conditional probabilities. The transformation from nucleotide sequences to k-mer histograms allows for fast, alignment-free, statistical measures to be used in comparisons.

The selection of this *k* parameter depends on the size of the input sequences. MeShClust automatically computes the *k* by first taking the *log*_4_ of the average sequence length, then by subtracting 1. We empirically found that this formula preserves enough information to accurately determine similarity. A smaller *k* value decreases the amount of memory needed for each histogram and the time required to calculate the alignment-free statistics by a factor of 4 for each nucleotide (26).

Once sequences are converted to k-mer histograms, the classifier is trained in the next stage.

### Identifying similar sequences

MeShClust utilizes a classifier to predict similar sequences to a query sequence. The similarity is determined according to an identity score obtained by global alignment (22, 23). With regard to a query sequence, similar sequences can be viewed as one class and dissimilar sequences as the other. Therefore, this task can be represented as a classification task. To this end, we used a GLM (27) for classifying these two classes. As a first step, MeShClust samples a roughly equal number of pairs of similar and dissimilar sequences based on a user-defined cutoff. A large number of pairs of sequences is needed to be sampled. Therefore, about 1500 sequence pairs are sampled. Similar sequences are labeled with 1’s and the dissimilar sequences with −1’s. After that, four features are extracted for each pair of sequences. These four features are selected according to a comprehensive evaluation of alignment-free k-mer statistics (26). The first feature is sequence length difference (Equation **3**) × Czekanowski similarity (Equation **4**). Length difference × Manhattanalgorithm, if applicable, is used for finding similar sequences to that center. After that, the mean shift is applied on alldistance^2^ (Equation **5**) represents the second feature. The thirdand the forth features are the Pearson coefficient (Equation **6**), and Kulczynski22 (Equation **7**) × Length difference^2^

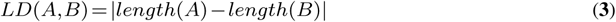

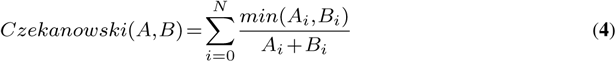

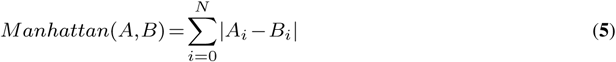

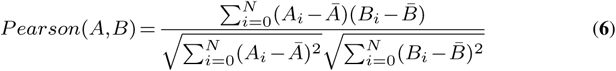

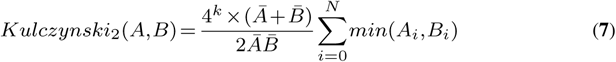

*A* and *B* are the two histograms representing two sequences; A*_i_* and B*_i_* are the counts of the *i^th^* k-mer in A and B; 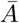 and 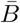 are the average counts of histograms A and B. Next, the four features are scaled between 0 and 1 and converted to similarity measures if necessary by subtracting the scaled value from 1. Then MeShClust utilizes an incremental, automatic process to train the GLM. Before training, the data is divided into two sets of equal sizes. Each set has roughly equal points representing similar and dissimilar sequence pairs. One set is used for training the classifier and the other is used for testing it. First, it trains the GLM using the first two features. If the accuracy calculated on the testing set is at least 97.5%, the training process is finished. Otherwise, it continues by adding another feature followed by evaluating the testing accuracy. This accuracy is measured by the average of the true positive rate (sensitivity) and the true negative rate (specificity). Once trained, the GLM is used for predicting sequences similar to a query sequence. We have observed that if the similarity is under 60%, it is not currently possible to accurately classify sequences using alignment free statistics, so alignment is used in those cases.

### Efficient data storage

Because the algorithm repeatedly selects and removes similar sequences to a query sequence, the largest time bottleneck was finding and removing multiple sequences from the input list. As a remedy, a data structure similar to a separate chaining hash table was implemented that allowed for faster retrieval and deletion of elements. This data structure consists of a list of smaller bins which specify a range of sequence lengths. Therefore, searching for similar sequences within a certain length range only affects a few bins when sequences are removed from the data structure. Using this data structure, initial centers can be found with relative ease, as discussed in the next section.

### Finding initial clusters

MeShClust aims at clustering the input sequences into distinct groups. Figure 1 diagrams this step. MeShClust gathers similar sequences into initial clusters. To start, input sequences are sorted based on length. The shortest sequence is the center of the first initial cluster. Then the classifier or the alignment sequences in the current cluster to calculate the updated mean. Next, the sequence closest to the new mean becomes the center of the cluster. Therefore, at each iteration of the algorithm, a better representative center of the cluster is found. This new center is used for the addition of more similar sequences to the cluster. This step is repeated until no similar sequences are left. At this point, the current cluster is set aside, and a new cluster is formed using the next closest sequence to the last center. The selection of the next center improves clustering by producing a semi-sorted list of sequences; neighboring clusters that may be merged later are grouped near each other. In effect, the combination of using a binary classifier and running the mean shift represents a “flat kernel”(15), except it only considers sequences not already placed in the initial clusters.

**Figure 1.**
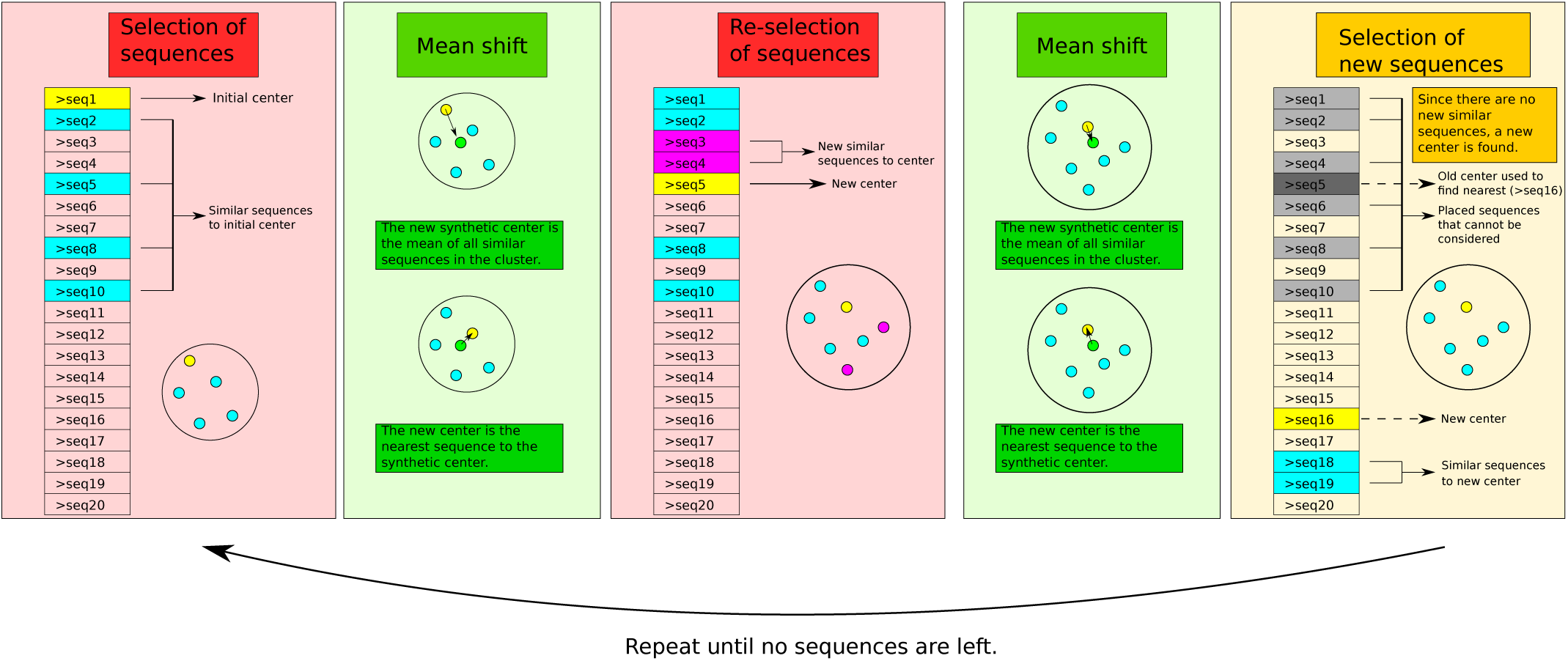
Overview of the initialization step of MeShClust, the first part of Algorithm 1. Before running this step the input sequences are sorted according to their lengths. The center of the first initial cluster is the shortest sequence in the input. The diagram shows the iterative process of finding initial clusters of sequences, getting better clusters from the selected sequences, and then finding the next closest cluster.

After grouping the sequences into initial clusters, the mean shift is used once again for forming the final clusters.

### Clustering

In the previous step, the classifier only considers the unplaced sequences; therefore, some of the initial clusters may have similar sequences in the already placed clusters. Further, the center of a cluster is updated in the initialization step by considering its sequences only. In this step, unlike the initial clustering, the mean shift considers sequences in neighboring clusters. These neighboring clusters include 5 clusters above and below the current cluster. Recall that these clusters are placed in a semi-sorted list. Centers that are close to each other, as determined by the classifier or by the alignment algorithm, are merged. If two centers are merged, the sequences that belong to each center are also merged.

## RESULTS

We start with defining multiple evaluation measures in order to evaluate MeShClust. These measures are intended to evaluate the quality of the predicted clusters as well as the time and the memory requirements of each software tool. After that, we discuss the results of comparing five sequence clustering software tools, including MeShClust.

### Evaluation criteria

We applied the following seven evaluation measures in evaluating MeShClust and four related tools: (i) intra-cluster similarity, (ii) inter-cluster similarity, (iii) silhouette score, (iv) purity, (v) normalized mutual information (NMI), (vi) time requirement, and (vii) memory usage. Only clusters of at least five sequences are considered in our analysis of the first three measures, except when it is indicated otherwise. The intra-cluster similarity is the average similarity between the sequence representing the center of a cluster and the other sequences in the same cluster. Sequence similarity is determined by calculating the identity score. To measure the dissimilarity between different clusters, we applied the inter-cluster similarity measure, which is the average similarity between different centers. Our third criterion is a variant of the silhouette score (28). This measure compares the suitability of placing a sequence in its current cluster to the suitability of placing this sequence in the closest cluster. We define *d_c_*(*s*) as the distance between the sequence *s* and the sequence representing its own cluster and *d_n_*(*s*) as the distance between *s* and the sequence representing the closest “neighboring” cluster. The distance is measured by subtracting the identity score from 100. Equation **8** defines the silhouette score.

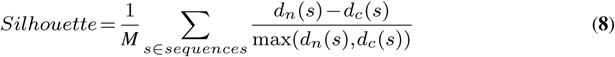

Here, *M* is the number of sequences in the set. The silhouette score ranges between 1 and −1; when the score is high, clusters are tighter and more separated from each other. A perceived problem with the silhouette score is that clusters consisting of a single element always have the value 1. Therefore, when the silhouette is very high, single element clusters may be inflating that value. A possible remedy for this issue is the redefinition of a cluster as a collection of at least five elements. However, a good silhouette value may not match actual “perfect” clusters. The silhouette is only a measure for combined separation and tightness, not correctness.

Our fourth and fifth evaluation measures do measure correctness. They are purity (Equation **9**) (29) and NMI (Equation **12**) (29), both of which are applied when the true clusters are known. These measures compare all clusters found, *F*, to the actual clusters, *A*, in a set of *N* sequences.

Purity (Equation **9**) measures how mixed each cluster is; if a predicted cluster only includes items from one real cluster, the purity is high. A disadvantage of this measure is that if every cluster is a single element, the purity will be 1.

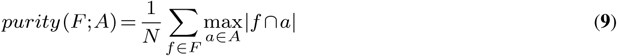

NMI corrects this by considering the probability that *F* and *A* contain the same data. To make these values comparable, this mutual information is normalized by the average entropy of *F* and of *A*. Equations **10, 11**, and **12** describe mutual information, entropy, and NMI.

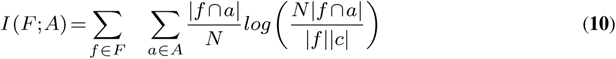

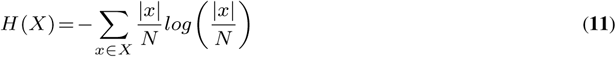

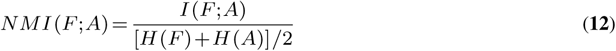

Next, we apply these evaluation measures to assessing the performance of MeShClust and four widely used tools.

### Comparison to related tools

We utilized both synthetic and real data sets in comparing the performance of MeShClust to the performances of UCLUST (12), CD-HIT (3, 11), DNACLUST (9), and wcdest (10, 30), which are widely used clustering tools. All related tools were ran using the default values for all parameters except the sequence identity parameter, for which we tried multiple values. However, wcdest does not have an identity parameter; the authors of wcdest provide an example parameter values equivalent to 96% identity, given as “-N 4 −l 100 -T 40 -H 72 --show clusters”. Therefore, we compared to wcdest only using the 96% identity when applicable. All other tools were also evaluated at 96% for comparison. Evaluations were done on a Dell OptiPlex 990 with 4 Intel i5-2500 processors running GNU/Linux (CentOS 7).

### Evaluation on synthetic data sets

As a start, we generated three data sets, which we call the 10%, the 25%, and the 100-centers sets (Supplementary Data Set 4–6). The three synthetic data sets were generated at 10%, 25%, and 25% mutation rates, respectively. Each of the 10% and the 25% data sets has 10 clusters, each of which consists of about 25 sequences. The 100-centers set has 100 clusters each of size 10. The length of a sequence is 1000 base pairs (bp) approximately.

To test the tolerance of the five tools to the identity parameter, we ran the tools using five identity scores (up to 10% above and below the actual identity score). Two of these identity scores are above the real identity score, and two are below it, and one approximately matches the identity score used for generating the clusters. These identities were selected to demonstrate to what degree a tool is tolerant to an inaccurate identity score. Table 1 shows the results of evaluating the four tools on the 10% data set using these identity scores: 75%, 80%, 85%, 90% and 95%. The true identity score is between 85% and 90%.

**Table 1.**
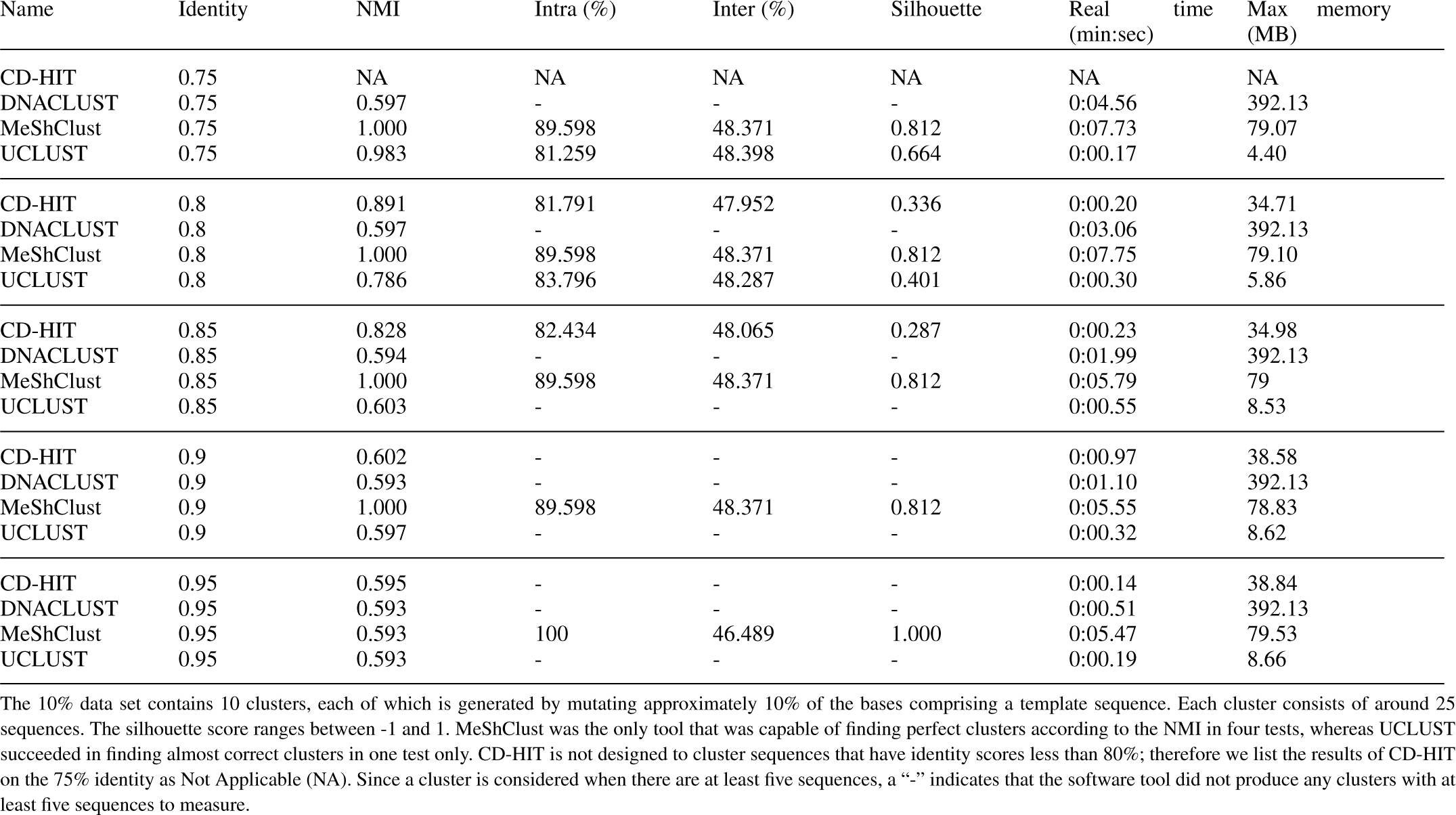
Comparison of the performances of MeShClust, UCLUST, CD-HIT, and DNACLUST on the 10%-mutation-rate synthetic data set.

When the true clusters are known, the performance is best measured by purity and NMI. On most synthetic data sets, the purity values achieved by all tools were 1, and the lowest value found was above 0.9. Thus NMI becomes the de-facto measure. On four out of the five tests, MeShClust obtained perfect NMI and purity scores. On those tests, MeShClust found perfect clusters, even though the identity parameter was inaccurate. Among the three related tools, UCLUST obtained almost perfect NMI score in one test only. CD-HIT and DNACLUST did not obtain perfect or close to perfect results on any of the five tests. Further, MeShClust achieves much better results in terms of the silhouette and the intra-cluster similarity scores. Compared to MeShClust’s intra-cluster score of 89.598 and silhouette of 0.812, the next highest of any tool on any identity was UCLUST with 83.796 similarity on the 80% identity and UCLUST with a silhouette of 0.664 on the 75% identity. Similar results were obtained on the 25% and the 100-centers data sets (see Supplementary Table 1). These results demonstrate that MeShClust is tolerant to inaccurate identity score. This tolerance is evident by the consistency of the high quality clusters obtained on different synthetic data sets at different identity levels.

### Evaluation on a comprehensive microbiome

Next, we aimed at evaluating the tools on real data; therefore, we obtained sequences from a microbiome study (31). We call this set the Costello set. About 1.1 million sequences comprise this set. Sequences in this set range between 200 and 400 base pairs (bp). Before evaluating the tools on the Costello set, we generated similar, smaller synthetic sets because the real clusters are unknown. We generated the 15K and the 150K sets consisting of 15 and 150 thousand sequences, respectively (Supplementary Data Sets 7 and 8). A synthetic cluster contains around 75 sequences, forming 200 and 2000 actual clusters, respectively. These clusters were generated using 3% mutation rate; however, the actual mutation is usually higher than 3% due to randomization.

As before, we evaluated the tools using the following identity scores: 83%, 87%, 90%, 93%, 96%, and 97% (Supplementary Table 2). On the 15K set, MeShClust obtained perfect or almost perfect NMI in six tests, demonstrating its ability to find the real clusters even when the sequence similarity parameter is inaccurate. CD-HIT obtained perfect or almost perfect NMI in four tests, whereas UCLUST obtained its best, 0.98 and 0.95 NMI, in two tests. DNACLUST performed well in one test only, its best NMI was 0.89. When the identity threshold is 96%, wcdest has 0.879 NMI, outperforming all other tools except MeShClust, which achieved 0.998 NMI. With respect to time required, MeShClust takes more time than UCLUST or CD-HIT, but MeShClust is faster than DNACLUST and wcdest. In terms of memory use, MeShClust takes more memory than UCLUST, CD-HIT, and wcdest, but it uses only around 90 MB, which is readily available on modern hardware. MeShClust has the highest silhouette scores except for a few cases where DNACLUST or wcdest has higher values. However, DNACLUST at 97% only had 0.2% of its clusters containing at least five sequences. Thus, DNACLUST’s silhouette of 0.988 is due to a small number of clusters because the majority of its clusters are too small. A similar phenomenon happens with wcdest.

**Table 2.**
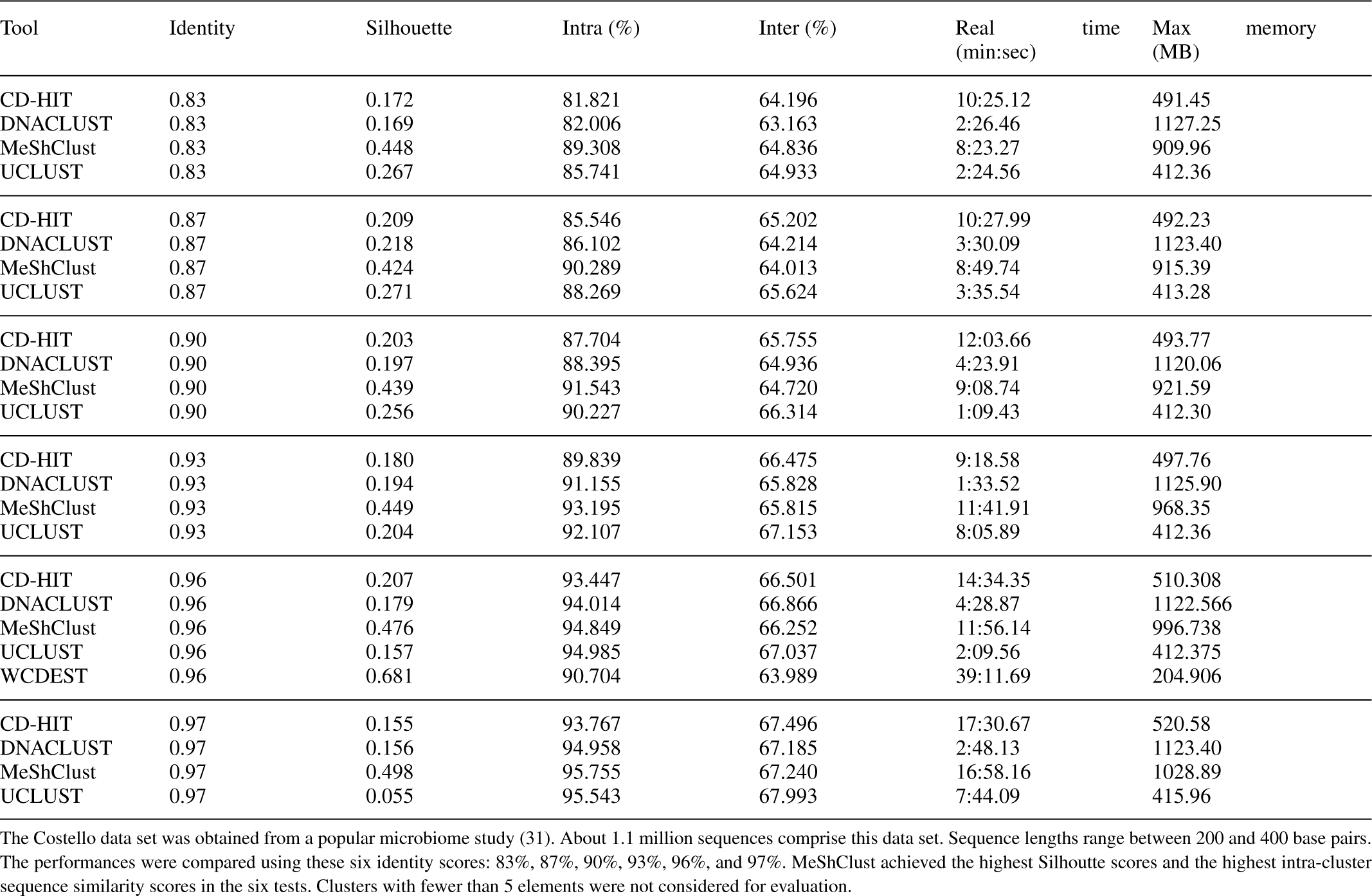
Comparison of the performances of the five tools on the Costello data.

MeShClust achieved similar, even better, results on the 150K data set (Supplementary Table 2). Specifically, it obtained an NMI of 1 in all but one test, where it has NMI of 0.984, outperforming all related tools on this data set. The second best performance was achieved by CD-HIT, which scored perfect or almost perfect NMI in four tests. The silhouette scores obtained by wcdest and MeShClust were comparable (0.94 versus 0.93). Even though wcdest had a slightly higher silhouette score, again wcdest only had 0.6% of its clusters with at least five sequences; therefore, its silhouette of 0.941 is due to 0.6% only of wcdest’s output. In contrast, MeShClust had *>* 95% of its clusters containing at least 5 sequences except on the 97% identity test. *Interestingly, the clusters produced by MeShClust were stable across all identity cutoffs demonstrated by almost the same values of NMI, silhouette, intra-clustering similarity, and the inter-clustering similarity.*

After that, we evaluated the five tools on the Costello set (Supplementary Table 2), which consists of 1.1 million real sequences. Table 2 shows these results. MeShClust obtained higher silhouette and intra-cluster similarity scores than the four related tools in the six tests. With respect to the time required, MeShClust took 8-17 minutes to cluster the Costello set. On average, MeShClust takes 20.4% longer time than the other tools, and around 6 minutes more than UCLUST. However, on computers with many CPU cores, this time gap will be lowered because of MeShClust’s parallel algorithm. Regarding the memory usage, MeShClust takes a manageable amount of memory of approximately 1GB, which is available on almost all personal computers. Concerning the quality of the clusters, MeShClust has the highest silhouette of any tool besides wcdest, averaging 0.1915 higher silhouette than the next highest silhouette on the Costello data set (excluding wcdest). However, the percent of clusters produced by wcdest at or above 5 sequences was 2.4%, implying that wcdest missed the majority of the real clusters. In contrast, all the other tools had at least 72 times as many clusters at or above 5 sequences as wcdest. *These results show that the clusters produced by MeShClust are more separable and more compact than those produced by the related tools.*

### Evaluation on viral genomes

Entire viral genomes (32) were used for testing MeShClust on much longer sequences, with sequence lengths averaging 6625 base pairs and with some sequences over 13000 base pairs. The data set contained 7 different families or genera, totaling 96 sequences (Table 3). Genomes from genera Badnavirus, Caulimovirus, Reptarenavirus, Soymovirus, and Spumavirus were used, and genomes from families Amalgaviridae and Birnaviridae were combined to create this data set. Since the taxonomic data was known, the real clusters were created by grouping every genera/family into a cluster. Therefore, purity and NMI values can be calculated for this test.

**Table 3.** The viral data set.

| Genome | Average length | Sequences in cluster |
| --- | --- | --- |
| Complete genome Badnavirus | 7574 | 42 |
| Segment B Birnaviridae | 2836 | 8 |
| Complete genome Spumavirus | 12307 | 6 |
| Complete genome Amalgaviridae | 3369 | 8 |
| Complete genome Caulimovirus | 7940 | 11 |
| Complete genome Soymovirus | 8088 | 5 |
| Segment S Reptarenavirus | 3400 | 4 |
| Segment A Birnaviridae | 3173 | 8 |
| Segment L Reptarenavirus | 6900 | 4 |

The viral data set consists of 9 clusters and 96 viral genomes. Viral genoms were obtained from viruSITE (32).

An analysis of the real clusters showed an intra-cluster similarity of 59% and an inter-cluster similarity of 40%. Therefore, identities 43%, 47%, 50%, 53%, and 57% were used for comparing MeShClust and UCLUST. These identities were too low for CD-HIT or DNACLUST; therefore they were not included in this test.

Table 4 shows the results on the viral set. MeShClust and UCLUST had very similar purity values over the viral data sets (0.002 average difference). However, MeShClust outperformed UCLUST on those tests via NMI by an average difference of 0.25 on a scale from 0 to 1, while using similar memory requirements and a manageable amount of time.

**Table 4.** MeShClust and UCLUST were used for clustering viral genomes

| Name | Id | Purity | NMI | Intra (%) | Inter (%) | Silhouette | Real time (min:sec) | Max memory (MB) |
| --- | --- | --- | --- | --- | --- | --- | --- | --- |
| MeShClust | 0.43 | 0.573 | 0.583 | 51.742 | 49.928 | 0.297 | 0:25.00 | 81.38 |
| UCLUST | 0.43 | 0.467 | 0 | 38.428 | 100 | 0.384 | 0:11.06 | 72.18 |
| MeShClust | 0.47 | 0.667 | 0.677 | 54.705 | 45.016 | 0.269 | 0:31.33 | 81.26 |
| UCLUST | 0.47 | 0.700 | 0.485 | 63.044 | 42.667 | 0.302 | 0:18.98 | 74.48 |
| MeShClust | 0.50 | 0.917 | 0.865 | 72.016 | 39.093 | 0.501 | 1:28.06 | 81.53 |
| UCLUST | 0.50 | 1 | 0.729 | 91.413 | 37.626 | 0.830 | 0:32.13 | 75.79 |
| MeShClust | 0.53 | 1 | 0.866 | 77.635 | 37.424 | 0.585 | 2:48.81 | 81.66 |
| UCLUST | 0.53 | 1 | 0.595 | 92.527 | 37.824 | 0.840 | 0:34.09 | 77.33 |
| MeShClust | 0.57 | 1 | 0.687 | 91.206 | 37.864 | 0.824 | 6:15.50 | 82.10 |
| UCLUST | 0.57 | 1 | 0.575 | 96.024 | 38.651 | 0.905 | 0:35.55 | 77.85 |
| Ground truth | - | 1 | 1 | 59.004 | 39.764 | 0.274 | - | - |

The viral data set containing 96 genomes was obtained from viruSITE (32). Since the clusters were so small, the intra, inter, and silhouette considered any cluster, not just clusters of at least 5 elements. The purity was not 1 all around, getting as low as 0.467. Overall, MeShClust had similar purity values (0.002 average difference); however, MeShClust had an NMI value on average 61% higher NMI values than UCLUST. On the 43% data set, UCLUST only found 1 cluster, explaining its 100% inter-cluster distance and 0 NMI.

On the 53% test, MeShClust found an NMI of 0.866 and a silhouette of 0.585, while UCLUST found an NMI of 0.595 and a silhouette of 0.840. The silhouette of the real clusters is 0.274. Thus, a higher silhouette score does not directly imply having more accurate clusters in this case. Using a cutoff of 53%, UCLUST found 55 clusters, each averaging 1.636 sequences, whereas MeShClust found 19 clusters, each averaging 5.053; this data set consists of 9 real clusters (Genomes of Reptarenavirus and Birdnavirus are broken up into two segments). These statistics can be found in Supplementary Table 3. Because UCLUST’s median cluster size is 1, many of its sequences will get a silhouette value of 1 and very high intra cluster values. Therefore, a higher silhouette score does not indicate better clusters in this case. However, MeShClust’s clusters overlap with the real clusters more than the ones produced by UCLUST, evident by higher NMI.

## DISCUSSION

### Added benefits of the classifier

Recall that when the identity cutoff is greater than 60%, MeShClust uses a classifier to predict the identity score from a combination of alignment-free k-mer statistics. To assess the added benefits of the classifier over the alignment algorithm, we evaluated the performances of the two versions of MeShClust on the 15K data set. As expected, the alignment-only version is 16 minutes (198 times) slower, on average, than the classifier-based version (Supplementary Table 4). Using only alignment does not give higher quality clusters; in no case does the alignment-only version have higher NMI values than the classifier-based version. These results demonstrate the remarkable time reduction due to the classifier.

### Related algorithms

DBSCAN (33) is a “density-based” clustering algorithm. Like the mean shift, it is based upon the principle that clusters consist of densely-packed points. Both the mean shift and DBSCAN can discover the number of clusters on their own. DBSCAN depends on two parameters, the minimum number of close points and the minimum distance between close points. In contrast, the mean shift algorithm depends on only one parameter, the cluster bandwidth, which is the distance among points in the same cluster. The average time of DBSCAN is *O* (*n*log(*n*)) when using an efficient data structure such as R* tree. However, this degrades into a quadratic time if an efficient data structure is not used. Similarly, the mean shift is a quadratic algorithm in theory because each point represents the center of an initial cluster. However, using a reasonable similarity threshold, our implementation should take *O* (*mn*), where m is the number of clusters in the data set. This run time may degenerate to quadratic time if the similarity threshold is too high. DBSCAN considers adding a new point to a cluster if it is close to a at least a minimum number of points in the cluster. This new point and the points close to it are added to the cluster. For this reason, DBSCAN can find clusters of many different shapes. Overall, the structure of the algorithm is similar in nature to MeShClust’s implementation of the mean shift, except MeShClust uses the average histogram as the new center; points that are close to the average are added to the cluster.

MCL (34) (35) is a graph based clustering algorithm. It uses random Markov walks to collect “flow” in a graph, that is, if a random walk is performed in a graph, most likely that walk will end in the same cluster as it started. Using an all-vs-all adjacency matrix, MCL finds clusters by converting the distances into transition probabilities, i.e. constructing a stochastic matrix, in which each column sums to 1. The core of this algorithm is a process involving squaring the stochastic matrix. An entry in the newly-squared matrix represents the probability of reaching a node from another directly or via one more node. In other words, squaring the matrix simulates a step along another edge. This process is performed until each element is either 0 or 1. Although it consists of matrix operations, efficient structures such as sparse matrices allow for fast computation. Using a BLAST-style input, the alignments can be pre-computed for reuse, and the parameters are determined by BLAST, not by the algorithm itself. In sum, MCL uses a different approach that is based on graph theory, whereas the mean shift is based on density-based optimization.

## CONCLUSION

DNA sequence clustering algorithms have many applications. Nonetheless, the widely applicable tools depend on greedy algorithms, which do not necessarily produce the best results. Further, the related tools are sensitive to the sequence similarity parameter provided by the user. Often times, the exact value of this parameter is not known, resulting in inaccurate clusters. Our clustering software, MeShClust, is a novel tool that utilizes the mean shift algorithm. *MeShClust is the first application of the mean shift to clustering DNA sequences and one of few applications of the mean shift algorithm in bioinformatics*. Unlike the related greedy approaches, the theory behind the mean shift guarantees convergence to local optimal points, resulting in higher quality clusters. Further, most sequence clustering tools use a slow quadratic algorithm for sequence alignment. In contrast, MeShClust reduces the dependency on alignment algorithms by using a novel, machine-learning-based, alignment-assisted method for computing sequence similarity. Furthermore, *this is the first attempt to formulate the task of identifying similar sequences as a classification task*. When tested on multiple synthetic and real data sets, MeShClust outperformed the related tools with a clear margin, advancing the methodology in the field of sequence analysis.

## SUPPLEMENTARY DATA

The C++ source code, the executables of MeShClust, and the synthetic data sets are available at NAR online as Supplementary Data Sets 1–9 and Supplementary Tables 1–4. MeShClust is also available at https://github.com/TulsaBioinformaticsToolsmith/MeShClust

## ACKNOWLEDGMENTS

We are thankful to the anonymous reviewers for taking the time to review this manuscript. Their comments and suggests have improved the software and the manuscript. We would like to thank Joseph Valencia and Robert Geraghty for their help with coding the GLM and the alignment algorithm.

